# Area-Based Selection of Binding Interfaces for Structural Prediction of Protein–Protein Complexes

**DOI:** 10.1101/2025.10.07.680849

**Authors:** Yong Xiao Yang, Bao Ting Zhu

## Abstract

Protein–protein interaction is a fundamental process in all biological systems, and the structural information of a protein–protein complex may provide important mechanistic details and insights into the biological processes involved. Elucidation of the rules underlying the interface specificity in protein–protein interactions is of great value for the correct prediction of the structures of a protein–protein complex. In the present study, we have developed the area-based methods for selecting near-native interfaces for protein–protein interactions. The quantitative relationship between different areas in the predicted structure of protein–protein complex and the predicted accuracy was explored using linear and nonlinear models. The predicted accuracy is characterized using the root mean square deviation (L_RMSD) of ligands. The performances of the newly-developed area-based models for selecting near-native interfaces for protein–protein binding interactions based on the partners’ structures at unbound or bound states are better than (or at least comparable to) those of the existing, more sophisticated method(s). The success rates of some models are above 90% (some are close to 100%), which indicates the importance and effectiveness of the area-based interface selection. The area-based methods presented in this work may shed lights on the final resolution of the interface selection problem in the field of protein–protein complex structure prediction and also on the rules of interface specificity for protein–protein interactions from a geometric area perspective. The principles developed this work also shed lights on understanding the protein-protein binding mechanisms from an area perspective.

**GRAPHICAL ABSTRACT:** 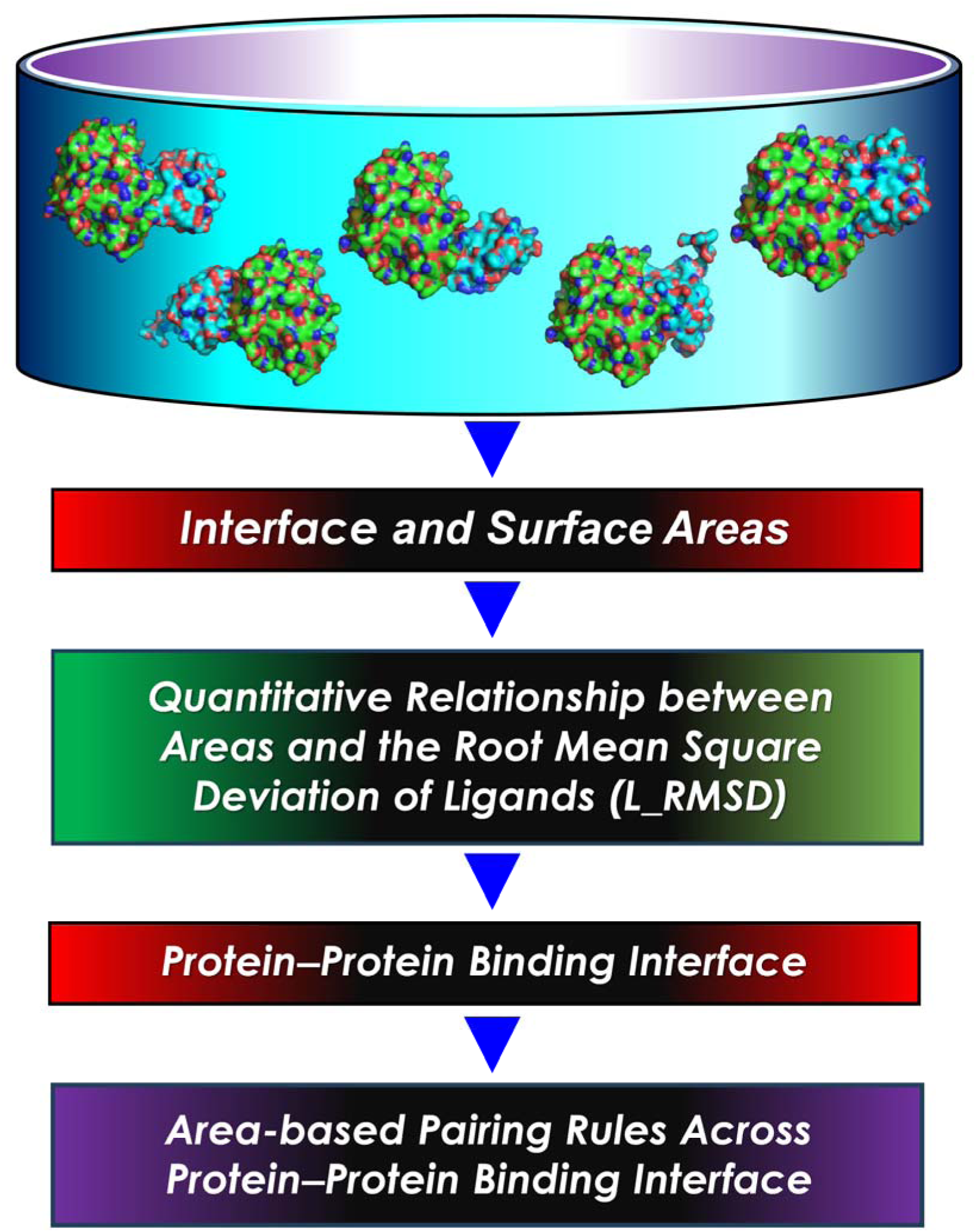

## 1. INTRODUCTION

In a living system, nearly all biological processes involve protein–protein interactions [1,2]. The structures of the protein–protein complexes are usually characteristic manifestations of the specific protein–protein interactions and the complexity of the biological processes involved. Experimental determination or theoretical prediction of the structures of the protein–protein complexes is among the most active research areas in biological, biophysical and computational sciences [3–5]. While the experimental methods to decipher protein–protein complexes are usually time-consuming and extremely labor intensive, the theoretical and computational methods are often limited by insufficient accuracy [5]. Hence, developing new and effective theoretical and computational methods with higher accuracy is still an area of research today with high importance and urgency.

Since the first work of using computers to generate possible protein–protein interaction models published in 1978 [6], many studies have been conducted for protein–protein docking which have become the main theoretical methods for the structural prediction of the protein–protein complexes [7–9]. The first step in protein–protein docking involves the generation of possible binding modes with geometrical feasibility and reasonability [9–11], and the second step is the selection of near-native docking decoys with effective scoring functions [12,13]. The reliable experimental or predicted binding site information can facilitate the structure prediction of the protein–protein complexes [14–20]. Sometimes, the structural prediction of a protein–protein complex fails at the second step, *i.e.*, the near-native structures do exist among the docking decoys, but are not successfully picked out. This failure might be partly due to the lack of a clear understanding of the rules that govern the interface specificity in protein–protein interactions, which is more complex compared to the base-pairing in the DNA structure [21]. Although different pairing potentials at the residue levels have been developed for structure prediction of a protein–protein complex [22–24], improvements in accuracy and effectiveness are still needed in interface selection for better prediction of the protein–protein interactions.

Area is an important metric in protein structure-related research. In 1986, Eisenberg and McLachlan developed a solvent accessible surface area-based model for calculating the protein’s atom solvation energy [25]. In the first work about proteinLprotein docking [7], an approximate total interface area was employed to evaluate the docked complexes. The total Voronoï interface area was also adopted as a parameter in a new proteinLprotein docking scoring function developed by Bernauer *et al.* [26]. The desolvation energy term was usually approximated based on the solvent accessible surface areas in the scoring functions for selecting near-native docking decoys [27,28].

In the present study, the interface and surface areas are divided into different parts based on the physicochemical properties of different amino acid types. These different kinds of interface and surface areas are employed to develop area-based methods for selecting near-native interfaces among a large number of docking decoys. The rules of interface specificity in protein–protein interactions are explored from a geometric area perspective.

## 2. MATERIALS AND METHODS

### 2.1. Experimental procedures for generating the datasets

The structures of protein–protein complexes used in this work were from PPI4DOCK [29,30] and protein–protein docking benchmark version 5 [31] and 5.5 [32]. After removing redundant structures using TM-align [33] and filtering based on several criteria specified below, 432 structures of the protein–protein complexes at the bound state and 116 structures at the unbound state were retained for analysis. The criteria included: (***i***) no more than 1500 residues in the structure of protein–protein complex; (***ii***) at least 25 residues in the structure of each binding partner; (***iii***) the number of chains in the structure at bound state is consistent with the structure at unbound state; (***iv***) the number of residues in the receptor structure at bound and unbound states is higher than or equal to the number of residues in the ligand structure; (***v***) both heavy and light chains exist in the antibody structure; (***vi***) the number of residues in the antibody structure is higher than 400; (***vii***) the number of residues in the antibody structure is higher than that in the structure of a protein antigen; (***viii***) there exist at least one near-native structure among the 2000 docking decoys generated by ZDOCK3.0.2 [34]. The near-native structures are the docking decoys with L_RMSD smaller than 5 Å. The L_RMSD is the root mean square deviation between the Cα atoms of ligands in docking decoy and the experimental structure after superimposition of the two corresponding receptor structures. The complexes are divided into 3 groups and 8 sets (**Table 1, Supplementary File 1**). The 3 groups are dimer group (SET-1, SET-2 and SET-3), multimer group (SET-4, SET-5 and SET-6), and antibody–protein antigen group (SET-7 and SET-8).

**Table 1.** Overview of the datasets used in this study.

| Group | SET No. | Number of complexes | Number of docking decoys | Number of near-native structures (L_RMSD < 5 Å) | Percentage of near-native structures (%) | State of the binding partners' structures | Source | References |
| --- | --- | --- | --- | --- | --- | --- | --- | --- |
| **Dimer group** | 1 | 190 | 379683 | 937 | 0.25 | bound | PPI4DOCK | [29] |
|  | 2 | 138 | 275751 | 719 | 0.26 | bound | Docking benchmark version 5 | [31] |
|  | 3 | 81 | 161891 | 340 | 0.21 | unbound |  | [31] |
| **Multimer group** | 4 | 19 | 37968 | 92 | 0.24 | bound | PPI4DOCK | [29] |
|  | 5 | 42 | 83943 | 157 | 0.19 | bound | Docking benchmark version 5 | [31] |
|  | 6 | 14 | 27971 | 30 | 0.11 | unbound |  | [31] |
| **Antibody–protein antigen group** | 7 | 43 | 85877 | 155 | 0.18 | bound | PPI4DOCK, Docking benchmark version 5 and 5.5 | [29,31,32] |
|  | 8 | 21 | 41943 | 58 | 0.14 | unbound | Docking benchmark version 5 and 5.5 | [31,32] |

To investigate the gaps between the structural prediction of a protein–protein complex and its affinity prediction, the affinity data were selected from 76 protein–protein complexes, which included 52 dimers and 24 multimers [35] obtained from the protein–protein binding affinity benchmark [31], and 33 antibody–protein antigen complexes [36] from the antibody–protein antigen binding affinity benchmark [32]. These selected complexes, in general, are considered to have more reliable experimental binding affinity data. After removing the repeated complexes and filtering based on the above two criteria (*i.e.*, at least 400 residues in the antibody structure; no inserted fragments in the sequence of the wild-type structure), 79 complexes were retained for analysis. These complexes were placed into one combined SET and divide into three subsets: SUBSET-1 (33 dimers), SUBSET-2 (9 general multimers) and SUBSET-3 (37 antibody–antigen complexes) (**Supplementary File 2**). Here, it should be noted that the affinity prediction is not the focus of this work, but the obtained results on affinity prediction are used to help illustrate the gaps between the interface selection (*i.e.*, structural prediction of a protein–protein complex) and the prediction of the affinity between a protein–protein complex.

### 2.2. Design of descriptors and calculation of the corresponding values

The rational design of accurate descriptors for the protein–protein complex structures is an important prerequisite for discovering the rules of interface specificity in protein–protein complexes. Geometric and energetic descriptors are among the frequently-used methods for further evaluating protein–protein interactions based on the structures. In the present study, only one class of geometric metric, *i.e.*, the area, is used to characterize the structures of protein–protein complexes.

As described in our previous works about affinity prediction [35,36], the surface and interface areas in the structure of a protein–protein complex were divided into 18 parts based on different types of amino acids. The 20 amino acid types are categorized into 4 groups: basic *AA*s (basic amino acids: HIS, ARG, LYS), nonpolar *AA*s (nonpolar, hydrophobic amino acids: ILE, PHE, LEU, TRP, ALA, MET, PRO, VAL), polar *AA*s (polar but uncharged amino acids: CYS, ASN, GLY, SER, GLN, TYR, THR), and acidic *AA*s (acidic amino acids: ASP, GLU). As summarized in **Table 2**, the surface areas are classified into 8 kinds: *RSA* (receptor surface area) of basic *AA*s (***A*_1_**), *RSA* of nonpolar *AA*s (***A*_2_**), *RSA* of polar *AA*s (***A*_3_**), *RSA* of acidic *AA*s (***A*_4_**), *LSA* (ligand surface area) of basic *AA*s (***A*_5_**), *LSA* of nonpolar *AA*s (***A*_6_**), *LSA* of polar *AA*s (***A*_7_**), and *LSA* of acidic *AA*s (***A*_8_**); the interface areas are divided into 10 kinds: basic *AA*s∼basic *AA*s (***A*_9_**), nonpolar *AA*s∼nonpolar *AA*s (***A*_10_**), polar *AA*s∼polar *AA*s (***A*_11_**), acidic *AA*s∼acidic *AA*s (***A*_12_**), basic *AA*s∼nonpolar *AA*s (***A*_13_**), basic *AA*s∼polar *AA*s (***A*_14_**), basic *AA*s∼acidic *AA*s (***A*_15_**), nonpolar *AA*s∼polar *AA*s (***A*_16_**), nonpolar *AA*s∼acidic *AA*s (***A*_17_**), polar *AA*s∼acidic *AA*s (***A*_18_**). Since only several hundred structures of the protein–protein complexes were used in this work, the surface and interface areas were not further divided into too many different parts and the 20 amino acid types were not considered separately, which might help avoid over-analysis.

**Table 2.** Different kinds of the surface and interface areas used in the analysis of the docking structures of the protein–protein complexes.

| Abbreviation | Region | Binding partner 1 | Binding partner 2 |
| --- | --- | --- | --- |
| $A_1$ | Solvent accessible surface | *RSA* (receptor surface area) of basic AAs (amino acids) | water molecules |
| $A_2$ | | *RSA* of nonpolar AAs | |
| $A_3$ | | *RSA* of polar AAs | |
| $A_4$ | | *RSA* of acidic AAs | |
| $A_5$ | | *LSA* (ligand surface area) of basic AAs | |
| $A_6$ | | *LSA* of nonpolar AAs | |
| $A_7$ | | *LSA* of polar AAs | |
| $A_8$ | | *LSA* of acidic AAs | |
| $A_9$ | Interface | basic AAs | basic AAs |
| $A_{10}$ | | nonpolar AAs | nonpolar AAs |
| $A_{11}$ | | polar AAs | polar AAs |
| $A_{12}$ | | acidic AAs | acidic AAs |
| $A_{13}$ | | basic AAs | nonpolar AAs |
| $A_{14}$ | | basic AAs | polar AAs |
| $A_{15}$ | | basic AAs | acidic AAs |
| $A_{16}$ | | nonpolar AAs | polar AAs |
| $A_{17}$ | | nonpolar AAs | acidic AAs |
| $A_{18}$ | | polar AAs | acidic AAs |

For convenience of calculation, the chain identifier of the receptor or ligand is renamed as A or B, respectively, regardless of whether the receptor (or ligand) is a monomer or multimer. The residues in the receptor or ligand were renumbered from 1. The interface residue contact areas and atom solvent accessible surface areas in the structure of a protein–protein complex were computed using Q_contact_ [37] and dr_sasa [38], respectively. Among all 1096000 ((432 + 116) × 2000) docking decoys, there were 973 structures for which the interface contact areas could not be normally calculated with Q_contact_ [37]. The reason might be due to the existence of atomic clashes. These structures were not used to train the models and evaluate the performances. There were 2488 near-native structures (L_RMSD < 5 Å, at least one for each complex) among the remaining 1095027 docking decoys. The percentages (%) of the near-native structures in the 8 sets ranged between 0.1 and 0.3 (**Table 1**).

### 2.3. Linear models for interface selection

In the present study, the simplest models, *i.e.*, linear empirical functions, were generated for the selection of the binding interfaces between a protein–protein complex and also for further analysis. In contrast to the more complicated machine learning methods, it was our intention that the possible meanings of the linear models with explicit formations might be readily explained in more clear terms, which, in turn, may offer insights into the underlying rules and principles.

The training sets for binding interface selection contained equal numbers of near-native structures (L_RMSD < 5 Å) and non-near-native decoys. All the near-native structures were included, and the non-near-native decoys were stochastically selected for each complex in the 8 sets shown in **Table 1**. The process was repeated 100 times. Additionally, the training sets were also composed of: ***1***) all decoys with L_RMSD < 5 Å; ***2***) all decoys with L_RMSD < 10 Å; or ***3***) all decoys with L_RMSD < 15 Å; or ***4***) all decoys with L_RMSD < 20 Å; or ***5***) all decoys. As a result, there are 840 subsets for training the linear empirical models. All different combinations of the 18 area-based descriptors were considered to select the near-native interfaces. Finally, there were 220200120 ((2^18^–1) × 840) linear models used for interface selection. The formation of the linear models for interface selection is shown below:

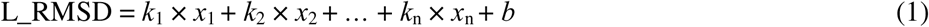

Here, *x*_1_, *x*_2_, …, *x*_n_ are the adopted area-based descriptors, *k*_1_, *k*_2_, …, *k*_n_ are the coefficients, and *b* is the constant term.

### 2.4. Neural network, random forest and mixed models for interface selection

To further explore the effectiveness of the area-based descriptors in interface selection, two frequently-used machine learning methods, *i.e.*, neural network [39,40] and random forest [41,42] were employed to train the selected models. The training sets were the same as those of the linear models for interface selection. In the neural network architecture, one or two layers were adopted, and the number of nodes were taken from 5, 10, 15 and 20. In the random forest architecture, the number of trees were chosen from 50, 100, 150 and 200, and the minimum number of observations per tree leaf was taken from 5, 10, 15 and 20. Accordingly, there were 20 kinds of neural network architectures and 16 kinds of random forest architectures. Because the stochasticity existed in the training process, 10 models were trained for each neural network or random forest architecture. Accordingly, there were a total of 168000 (840 × 20 × 10) neural network (NN) models and 134400 (840 × 16 × 10) random forest (RF) models.

Based on the neural network or random forest models, the mixed models, which are the linear combinations (based on linear regression) of the neural network or random forest models, were generated to further improve the performances in interface selection. The training sets were assigned as those for the linear, neural network and random forest models. There were 13761720 ((2^14^ – 1) × 840) mixed models based on 14 representative neural network models and 3439800 ((2^12^–1) × 840) mixed models based on 12 representative random forest models.

### 2.5. Previous models for near-native interface or structure selection and affinity prediction

According to previous studies [13,30], the scoring function adopted by ZDOCK3.0.2 is one of the best empirical models for selecting the near-native structures of a protein–protein complex, which employs geometric shape complementarity, electrostatic energy and knowledge-based pair potentials [23,34]. The results of this effective scoring function are compared with the results produced by models developed in this study.

In our previous works, there were 60 models for protein–protein binding affinity prediction [35] and 37 models for antibody–protein antigen binding affinity prediction [36]. The 97 models were also used for interface selection of protein–protein and antibody–protein antigen complexes.

### 2.6. Metrics for evaluating the performances of the models

The performances of the structural prediction of the protein–protein complexes using docking and scoring functions are usually evaluated according to the success rates, *i.e.*, the percentage of positive complexes in a given set when the top *N* decoys are retained for each complex [12,13]. The positive complexes are those with near-native structures (L_RMSD < 5 Å) among the retained top *N* decoys ranked by the scoring function. In this work, the areas under the success rate curves are also employed to evaluated the performances of linear models.

The performance of affinity prediction is frequently evaluated using the Pearson’s correlation coefficient (*R*) [35,36,43], which is expressed as follows:

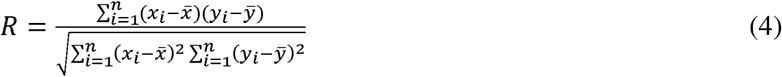

Here, *n* is the number of protein–protein complexes in a given set, *x_i_* (or *y_i_*) is the experimental (or predicted) binding affinity of the *i*^th^ protein–protein complex, *x̄* and *ȳ* are the average of the experimental and predicted binding affinities, respectively.

## 3. RESULTS

The flowchart of this work is shown in **Fig. 1**. The main steps include: (***i***) filtering of the original data; (***ii***) protein–protein docking to generate different binding poses. (***iii***) Calculation of the descriptors and training of the models. Different from the previous study on protein–protein binding interface selection [13], in this study the interface and surface areas are the only two parameters that are considered jointly for a given protein–protein complex. The interface and surface areas are divided into 18 parts as described in our recent studies on protein–protein binding affinity prediction [35,36]. (***iv***) Training of various models for interface selection, which include the linear models, neural network (NN) models, random forest (RF) models, NN-based models, and RF-based mixed models. Lastly, based on their performances, 63 representative models are selected for the selection of interfaces. The representative models include 3 linear models, 14 neural network (NN) models, 12 random forest (RF) models, 17 NN-based mixed models, and 17 RF-based mixed models. Here, it should be noted that in this study, selecting the “interfaces” or “decoys” often refers to the same process. For the two given protein binding partners with known structures, determining or selecting the binding interfaces is key to obtaining the correct structures of their complexes even without giving adequate consideration of their flexible conformational changes. Accordingly, selecting the correct binding interfaces implies selecting the correct docking decoys, *i.e.*, the near-native complex conformations.

**Figure 1.**
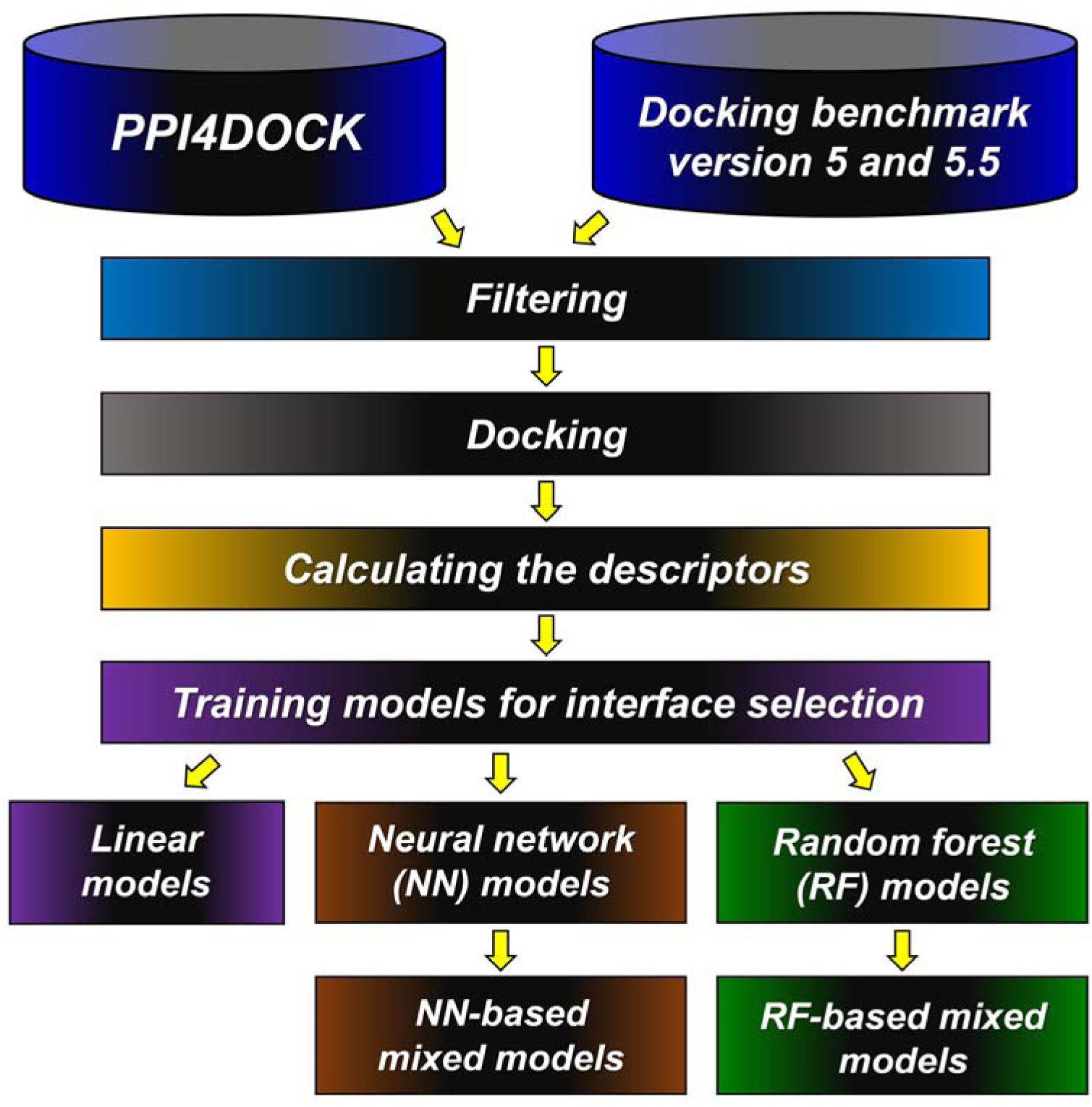
The flowchart of the binding interface selection for protein-protein interactions. The steps include: (***i***) filtering the original data; (***ii***) docking for genrating different possible binding poses; (***iii***) calculating the area-based desriptors; (***iv***) training various models for binding interface selection. The models inculde linear models, neural network (NN) models, random forest (RF) models, NN-based mixed models, and RF-based mixed models.

### 3.1. Representative area-based linear models for interface selection

The formations for the 3 representative models for interface selection are shown in **Table 3** (models 1–3, respectively). The shared descriptors of models 1–3 are ***A*_5_** (*LSA* of basic *AA*s), ***A*_10_** (nonpolar *AA*s∼nonpolar *AA*s), ***A*_12_** (acidic *AA*s∼acidic *AA*s), ***A*_14_** (basic *AA*s∼polar *AA*s), ***A*_15_** (basic *AA*s∼acidic *AA*s), ***A*_17_** (nonpolar *AA*s∼acidic *AA*s) and ***A*_18_** (polar *AA*s∼acidic *AA*s). The descriptors may play a more important role in the selection of binding interfaces.

**Table 3.** Formations of the representative linear models for interface selection.

| Model name and No. | Targeted application for interface selection | Formations |
| --- | --- | --- |
| **Linear model 1** | Dimer | $\begin{aligned} L\_RMSD = & -0.00179191603070685*A_5 + 0.00787201418786127*A_6 + 0.0794584161286932*A_9 - 0.0760255328665292*A_{10} - \\ & 0.0253270135617005*A_{11} + 0.101244860463670*A_{12} - 0.0414175112495662*A_{14} - 0.104891101254004*A_{15} - \\ & 0.0473629528979797*A_{16} + 0.0389434446932667*A_{17} - 0.0147033814705164*A_{18} + 37.9173495914677 \end{aligned}$ |
| **Linear model 2** | Multimer | $\begin{aligned} L\_RMSD = & 0.000465421504240589*A_1 + 0.0153037773426258*A_5 - 0.00486306286732690*A_7 - 0.00460698491611299*A_8 - \\ & 0.170358562365283*A_{10} + 0.0549390029184276*A_{11} + 0.428140887238351*A_{12} - 0.0934558885298738*A_{14} - \\ & 0.0789215506179263*A_{15} - 0.0685725890769897*A_{16} + 0.105466617292539*A_{17} - 0.0558538613554891*A_{18} + 49.7109738140069 \end{aligned}$ |
| **Linear model 3** | Ab–Ag complex | $\begin{aligned} L\_RMSD = & -0.00303649487993474*A_3 + 0.00614285429086451*A_4 + 0.00826466541141648*A_5 - 0.000500561405067450*A_7 + \\ & 0.00406983871857476*A_8 - 0.0241047212498659*A_9 - 0.0900757438416372*A_{10} + 0.0974560745669008*A_{12} - \\ & 0.0480681872044070*A_{13} - 0.0299927022381200*A_{14} - 0.0814104160696181*A_{15} + 0.145198666045445*A_{17} - \\ & 0.127039403354824*A_{18} + 33.6131307685756 \end{aligned}$ |

Note: $A_1, A_2, \dots, A_{18}$ are the 18 different parts of the surface and interface areas. L\_RMSD is the root mean square deviation of ligands after superimposition of the receptors. The Ab–Ag complexes refer to the antibody–protein antigen complexes.

The performances of ZDOCK3.0.2 [23,34] and the representative area-based linear models in different sets are shown in **Fig. 2** (stored in **Supplementary File 3**). The decoys in SET-3, SET-6 and SET-8 and those in SET-1, SET-2, SET-4, SET-5 and SET-7 were generated based on the partners’ structures at unbound and bound state, respectively. The performances of liner model 1 for dimers in SET-3 (success rate 33.33%) or linear model 2 for multimers in SET-6 (success rate 35.71%) are better than those of ZDOCK3.0.2 (success rate 27.16% in *SET-3* and 28.57% in SET-6), and the success rates (%) of these area-based linear models in SET-1, SET-2, SET-4, SET-5, SET-7 and SET-8 are lower than those of ZDOCK3.0.2.

**Figure 2.**
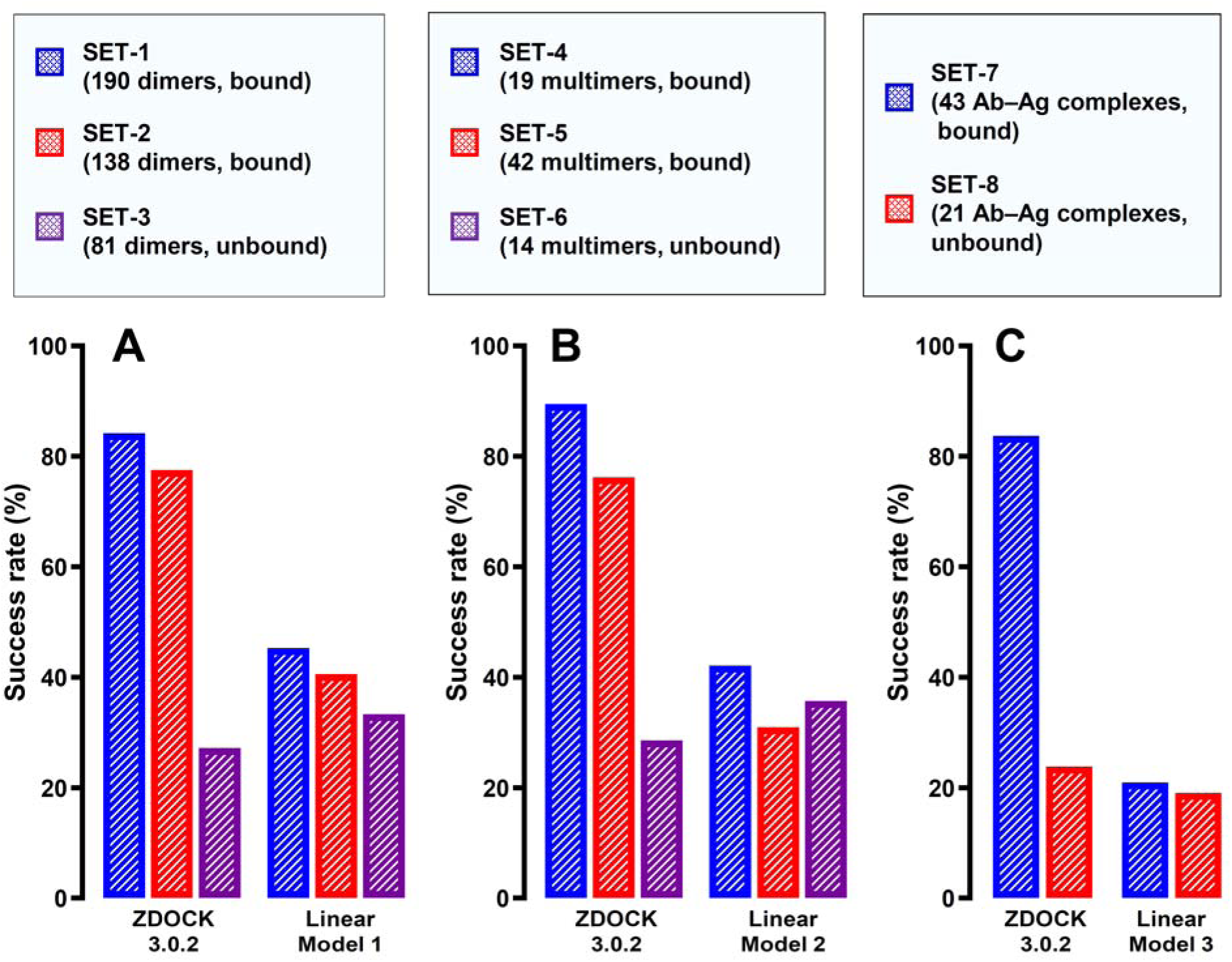
The performances of ZDOCK3.0.2 and 3 representative area-based linear models for protein–protein binding interface selection. **A.** The performances of ZDOCK3.0.2 and the linear model 1 (**Table 3**) in the dimer group. **B.** The performances of ZDOCK3.0.2 and the linear model 2 (**Table 3**) in the multimer group. **C.** The performances of ZDOCK3.0.2 and the linear model 3 (**Table 3**) in the antibody-protein antigen group. Note that there are 190 dimers in SET-1, 138 dimers in SET-2, 81 dimers in SET-3, 19 multimers in SET-4, 42 multimers in SET-5, 14 multimers in SET-6, 43 antibody-protein antigen complexes in SET-7, and 21 antibody-protein antigen complexes in SET-8, respectively. The possible interfaces for the complexes in SET-1, SET-2, SET-4, SET-5 and SET-7 and those in SET-3, SET-6 and SET-8 were separately generated from the partners’ structures at bound and unbound states, respectively. The performances shown in the figures are represented by the success rate (%) when the top 10 decoys are retained. The Ab–Ag complexes refer to the antibody–protein antigen complexes.

The success rates (%) of ZDOCK3.0.2 in SET-1, SET-2, SET-4, SET-5 and SET-7 (>75%) are much higher than those in SET-3, SET-6 and SET-8 (<30%). The scoring function of ZDOCK3.0.2 (composed of geometric shape complementarity, electrostatic energy and knowledge-based pair potentials) [23,34] is apt at selecting the near-native interfaces from the docking decoys generated based on partners’ structures at bound state. The gaps between the performances of area-based models for the two situations (bound and unbound) are not as big as that of the scoring function of ZDOCK3.0.2. The area-based models are relatively robust for docking from structures at bound or unbound states. These results indicate the effectiveness of these elementary descriptors, *i.e.*, interface and surface areas, in the interface selection for protein–protein interactions.

### 3.2. Representative area-based neural network, random forest and mixed models for interface selection

Based on the performances, some representative models were selected to exemplify the predictive power of the area-based methods in interface selection. These models are composed of 14 neural network (NN) models (**Table 4**), 12 random forest (RF) models (**Table 5**), 17 NN-based mixed models (**Table 6**) and 17 RF-based mixed models (**Table 7**).

**Table 4.** Performances of the 14 representative neural network models for interface selection.

| Model name and No. | Targeted application for interface selection | Success rate (%) (top 10 decoys, L_RMSD < 5 Å) |  |  |  |  |  |  |  |
| --- | --- | --- | --- | --- | --- | --- | --- | --- | --- |
|  |  | SET-1 (190 dimers, bound) | SET-2 (138 dimers, bound) | SET-3 (81 dimers, unbound) | SET-4 (19 multimers, bound) | SET-5 (42 multimers, bound) | SET-6 (14 multimers, unbound) | SET-7 (43 Ab–Ag complexes, bound) | SET-8 (21 Ab–Ag complexes, unbound) |
| NN model 1 | Dimer | **59.47** | 14.49 | 7.41 | 26.32 | 4.76 | 0.00 | 4.65 | 0.00 |
| NN model 2 |  | 12.63 | **57.25** | 24.69 | 21.05 | 16.67 | 0.00 | 0.00 | 0.00 |
| NN model 3 |  | 10.53 | 22.46 | **44.44** | 21.05 | 2.38 | 0.00 | 2.33 | 4.76 |
| NN model 4 |  | **36.32** | **36.23** | **33.33** | 36.84 | 19.05 | 7.14 | 4.65 | 4.76 |
| NN model 5 |  | **44.21** | **38.41** | **30.86** | 31.58 | 14.29 | 7.14 | 11.63 | 0.00 |
| NN model 6 | Multimer | 11.05 | 7.25 | 6.17 | **84.21** | 7.14 | 14.29 | 9.30 | 4.76 |
| NN model 7 |  | 15.79 | 17.39 | 8.64 | 5.26 | **59.52** | 7.14 | 4.65 | 0.00 |
| NN model 8 |  | 5.26 | 2.90 | 6.17 | 10.53 | 7.14 | **64.29** | 2.33 | 0.00 |
| NN model 9 |  | 23.68 | 15.94 | 9.88 | **36.84** | **47.62** | **28.57** | 6.98 | 4.76 |
| NN model 10 |  | 28.95 | 18.12 | 9.88 | **42.11** | **28.57** | **28.57** | 4.65 | 4.76 |
| NN model 11 | Ab–Ag complex | 2.63 | 3.62 | 3.70 | 0.00 | 7.14 | 0.00 | **65.12** | 23.81 |
| NN model 12 |  | 12.63 | 4.35 | 1.23 | 10.53 | 0.00 | 0.00 | 13.95 | **61.90** |
| NN model 13 |  | 2.11 | 7.25 | 7.41 | 5.26 | 9.52 | 0.00 | **46.51** | **38.10** |
| NN model 14 |  | 2.63 | 5.07 | 2.47 | 0.00 | 4.76 | 0.00 | **51.16** | **33.33** |

Note: The Ab–Ag complexes refer to the antibody–protein antigen complexes.

**Table 5.** Performances of the 12 representative random forest models for interface selection.

| Model name and No. | Targeted application for interface selection | Success rate (%) (top 10 decoys, L_RMSD < 5 Å) |  |  |  |  |  |  |  |
| --- | --- | --- | --- | --- | --- | --- | --- | --- | --- |
|  |  | SET-1 (190 dimers, bound) | SET-2 (138 dimers, bound) | SET-3 (81 dimers, unbound) | SET-4 (19 multimers, bound) | SET-5 (42 multimers, bound) | SET-6 (14 multimers, unbound) | SET-7 (43 Ab–Ag complexes, bound) | SET-8 (21 Ab–Ag complexes, unbound) |
| **RF model 1** | Dimer | **100** | 35.51 | 22.22 | 31.58 | 21.43 | 14.29 | 6.98 | 0.00 |
| **RF model 2** |  | 32.63 | **100** | 50.62 | 36.84 | 19.05 | 7.14 | 6.98 | 4.76 |
| **RF model 3** |  | 30.53 | 43.48 | **100** | 15.79 | 19.05 | 7.14 | 6.98 | 4.76 |
| **RF model 4** |  | **34.21** | **39.86** | **90.12** | 26.32 | 14.29 | 7.14 | 0.00 | 4.76 |
| **RF model 5** |  | **35.79** | **99.28** | **45.68** | 26.32 | 21.43 | 14.29 | 6.98 | 0.00 |
| **RF model 6** | Multimer | 33.68 | 25.36 | 11.11 | **100** | 19.05 | 21.43 | 6.98 | 0.00 |
| **RF model 7** |  | 24.74 | 24.64 | 8.64 | 21.05 | **100** | 42.86 | 6.98 | 4.76 |
| **RF model 8** |  | 12.63 | 10.14 | 7.41 | 31.58 | 23.81 | **100** | 2.33 | 4.76 |
| **RF model 9** |  | 29.47 | 26.81 | 9.88 | **36.84** | **97.62** | **50.00** | 4.65 | 4.76 |
| **RF model 10** |  | 34.21 | 28.26 | 13.58 | **42.11** | **78.57** | **42.86** | 4.65 | 4.76 |
| **RF model 11** | Ab–Ag complex | 17.37 | 16.67 | 9.88 | 31.58 | 14.29 | 7.14 | **100** | **47.62** |
| **RF model 12** |  | 12.11 | 7.25 | 2.47 | 21.05 | 9.52 | 0.00 | **32.56** | **100** |

Note: The Ab–Ag complexes refer to the antibody–protein antigen complexes.

**Table 6.** Performances of the 17 representative NN-based mixed models for interface selection.

| Model name and No. | Targeted application for interface selection | Success rate (%) (top 10 decoys, L_RMSD < 5 Å) |  |  |  |  |  |  |  |
| --- | --- | --- | --- | --- | --- | --- | --- | --- | --- |
|  |  | SET-1 (190 dimers, bound) | SET-2 (138 dimers, bound) | SET-3 (81 dimers, unbound) | SET-4 (19 multimers, bound) | SET-5 (42 multimers, bound) | SET-6 (14 multimers, unbound) | SET-7 (43 Ab–Ag complexes, bound) | SET-8 (21 Ab–Ag complexes, unbound) |
| NN-based mixed model 1 | Dimer | **63.16** | 20.29 | 12.35 | 26.32 | 11.90 | 7.14 | 4.65 | 0.00 |
| NN-based mixed model 2 |  | 23.68 | **63.04** | 32.10 | 26.32 | 19.05 | 0.00 | 0.00 | 4.76 |
| NN-based mixed model 3 |  | 23.68 | 35.51 | **51.85** | 21.05 | 4.76 | 0.00 | 4.65 | 0.00 |
| NN-based mixed model 4 |  | 16.32 | 42.75 | **51.85** | 21.05 | 11.90 | 0.00 | 2.33 | 0.00 |
| NN-based mixed model 5 |  | **44.21** | **47.83** | **43.21** | 31.58 | 19.05 | 0.00 | 16.28 | 4.76 |
| NN-based mixed model 6 |  | **45.79** | **44.93** | **43.21** | 36.84 | 19.05 | 7.14 | 2.33 | 0.00 |
| NN-based mixed model 7 | Multimer | 18.42 | 19.57 | 9.88 | **89.47** | 28.57 | 21.43 | 6.98 | 0.00 |
| NN-based mixed model 8 |  | 16.84 | 11.59 | 8.64 | 15.79 | **71.43** | 21.43 | 0.00 | 0.00 |
| NN-based mixed model 9 |  | 9.47 | 4.35 | 4.94 | 15.79 | 26.19 | **85.71** | 0.00 | 0.00 |
| NN-based mixed model 10 |  | 20.53 | 19.57 | 11.11 | **42.11** | **64.29** | **42.86** | 0.00 | 0.00 |
| NN-based mixed model 11 |  | 15.79 | 15.22 | 11.11 | **42.11** | **54.76** | **64.29** | 6.98 | 0.00 |
| NN-based mixed model 12 |  | 16.84 | 15.22 | 6.17 | **52.63** | **40.48** | **57.14** | 4.65 | 0.00 |
| NN-based mixed model 13 | Ab–Ag complex | 2.63 | 3.62 | 4.94 | 5.26 | 4.76 | 0.00 | **81.40** | 38.10 |
| NN-based mixed model 14 |  | 6.84 | 3.62 | 4.94 | 10.53 | 0.00 | 0.00 | 34.88 | **76.19** |
| NN-based mixed model 15 |  | 3.68 | 5.80 | 8.64 | 5.26 | 4.76 | 0.00 | **72.09** | **57.14** |
| NN-based mixed model 16 |  | 4.21 | 7.25 | 4.94 | 0.00 | 2.38 | 0.00 | **65.12** | **61.90** |
| NN-based mixed model 17 |  | 22.11 | 7.97 | 13.58 | 10.53 | 4.76 | 0.00 | **55.81** | **66.67** |

Note: The Ab–Ag complexes refer to the antibody–protein antigen complexes.

**Table 7.** Performances of the 17 representative RF-based mixed models for interface selection.

| Model name and No. | Targeted application for interface selection | Success rate (%) (top 10 decoys, L_RMSD < 5 Å) |  |  |  |  |  |  |  |
| --- | --- | --- | --- | --- | --- | --- | --- | --- | --- |
|  |  | SET-1<br>(190 dimers, bound) | SET-2<br>(138 dimers, bound) | SET-3<br>(81 dimers, unbound) | SET-4<br>(19 multimers, bound) | SET-5<br>(42 multimers, bound) | SET-6<br>(14 multimers, unbound) | SET-7<br>(43 Ab–Ag complexes, bound) | SET-8<br>(21 Ab–Ag complexes, unbound) |
| RF-based mixed model 1 | Dimer | **100** | 68.84 | 77.78 | 31.58 | 26.19 | 7.14 | 4.65 | 0.00 |
| RF-based mixed model 2 |  | **100** | 78.99 | 67.90 | 21.05 | 9.52 | 7.14 | 32.56 | 0.00 |
| RF-based mixed model 3 |  | 91.58 | **100** | 88.89 | 26.32 | 2.38 | 0.00 | 6.98 | 0.00 |
| RF-based mixed model 4 |  | 71.05 | **100** | 96.30 | 5.26 | 9.52 | 71.43 | 0.00 | 0.00 |
| RF-based mixed model 5 |  | 90.53 | 84.06 | **100** | 5.26 | 0.00 | 0.00 | 9.30 | 0.00 |
| RF-based mixed model 6 |  | 87.37 | 91.30 | **100** | 0.00 | 9.52 | 0.00 | 11.63 | 4.76 |
| RF-based mixed model 7 |  | **94.21** | **98.55** | **95.06** | 26.32 | 9.52 | 7.14 | 9.30 | 0.00 |
| RF-based mixed model 8 | Multimer | 25.79 | 47.10 | 2.47 | **100** | 76.19 | 35.71 | 4.65 | 0.00 |
| RF-based mixed model 9 |  | 42.11 | 7.25 | 1.23 | **100** | 33.33 | 42.86 | 6.98 | 0.00 |
| RF-based mixed model 10 |  | 40.00 | 13.04 | 1.23 | 78.95 | **100** | 71.43 | 2.33 | 0.00 |
| RF-based mixed model 11 |  | 12.63 | 6.52 | 0.00 | 57.89 | **100** | 92.86 | 2.33 | 0.00 |
| RF-based mixed model 12 |  | 17.37 | 0.00 | 3.70 | 63.16 | 57.14 | **100** | 4.65 | 4.76 |
| RF-based mixed model 13 |  | 17.37 | 7.25 | 0.00 | **84.21** | **90.48** | **85.71** | 0.00 | 0.00 |
| RF-based mixed model 14 | Ab–Ag complex | 18.95 | 15.94 | 9.88 | 15.79 | 14.29 | 7.14 | **100** | 61.90 |
| RF-based mixed model 15 |  | 1.58 | 2.17 | 3.70 | 0.00 | 2.38 | 0.00 | 44.19 | **100** |
| RF-based mixed model 16 |  | 17.37 | 10.14 | 7.41 | 15.79 | 9.52 | 0.00 | **76.74** | **85.71** |
| RF-based mixed model 17 |  | 16.32 | 0.00 | 1.23 | 84.21 | 16.67 | 0.00 | **93.02** | **76.19** |

Note: The Ab–Ag complexes refer to the antibody–protein antigen complexes.

As shown in **Tables 4** and **5**, the highest success rates for the top 10 decoys (L_RMSD < 5 Å) of the neural network (NN) models are >44% in all the 8 sets. Those of the random forest (RF) models are 100% for each set. The success rates of NN models *vs.* RF models can reach >30% *vs.* >35% for all the 3 sets in the dimer group, >25% *vs.* >40% for all the 3 sets in the multimer group, and >30% *vs.* >45% for all the 2 sets in the antibody–protein antigen group. The RF models are superior to the NN models, and the results of NN and RF models are better than those of the linear models.

As shown in **Tables 4** and **6**, the highest success rates for the top 10 decoys (L_RMSD < 5 Å) are increased by about 10% on average when the NN models are converted to NN-based mixed models; and those of the mixed models based on random forest models are still 100% for each set. The success rates of the NN-based mixed models *vs.* NN models can reach >40% *vs.* >30% for all the 3 sets in the dimer group, >40% *vs.* >25% for all the 3 sets in the multimer group, and >60% *vs.* >30% for all the 2 sets in the antibody–protein antigen group. As summarized in **Tables 5** and **7**, the success rates of the RF-based mixed models *vs.* RF models are >90% *vs.* >35% for all the 3 sets in the dimer group, >80% *vs.* >40% for all the 3 sets in multimer group, and >75% *vs.* >45% for all the 2 sets in the antibody–protein antigen group. Overall, the performances for several NN-based and RF-based mixed models are improved, but the success rates of the NN-based mixed models are still lower than the success rates of the RF-based mixed models.

In terms of the top-ranked 10 decoys, the performances of the best mixed models (*i.e.*, the RF-based mixed models 7, 13, 16 and 17 in **Table 7**) are much better than those of ZDOCK3.0.2 [23,34] when the docking decoys are generated from the partners’ structures at the unbound states (SET-3, SET-6 and SET-8), and are comparable to those of ZDOCK3.0.2 [23,34] in the three groups when the docking decoys are generated from the partners’ structures at the bound states (SET-1, SET-2, SET-4, SET-5 and SET-7). As shown in **Fig. 3**, the success rates (%) based on the top-ranked 100 decoys of the best RF-based mixed models are slightly higher than those of ZDOCK3.0.2 when the docking is based on the partners’ structures at the bound states, and are much higher than those of ZDOCK3.0.2 when the docking is based on the structures from unbound states. The expected performances of the random selections are also shown in **Fig. 3**. It is apparent that random selection is not as good as ZDOCK3.0.2 or the representative RF-based mixed models. These results demonstrate the effectiveness and superiority of the new prediction models based on interface and surface areas coupled with machine learning for interface selection.

**Figure 3.**
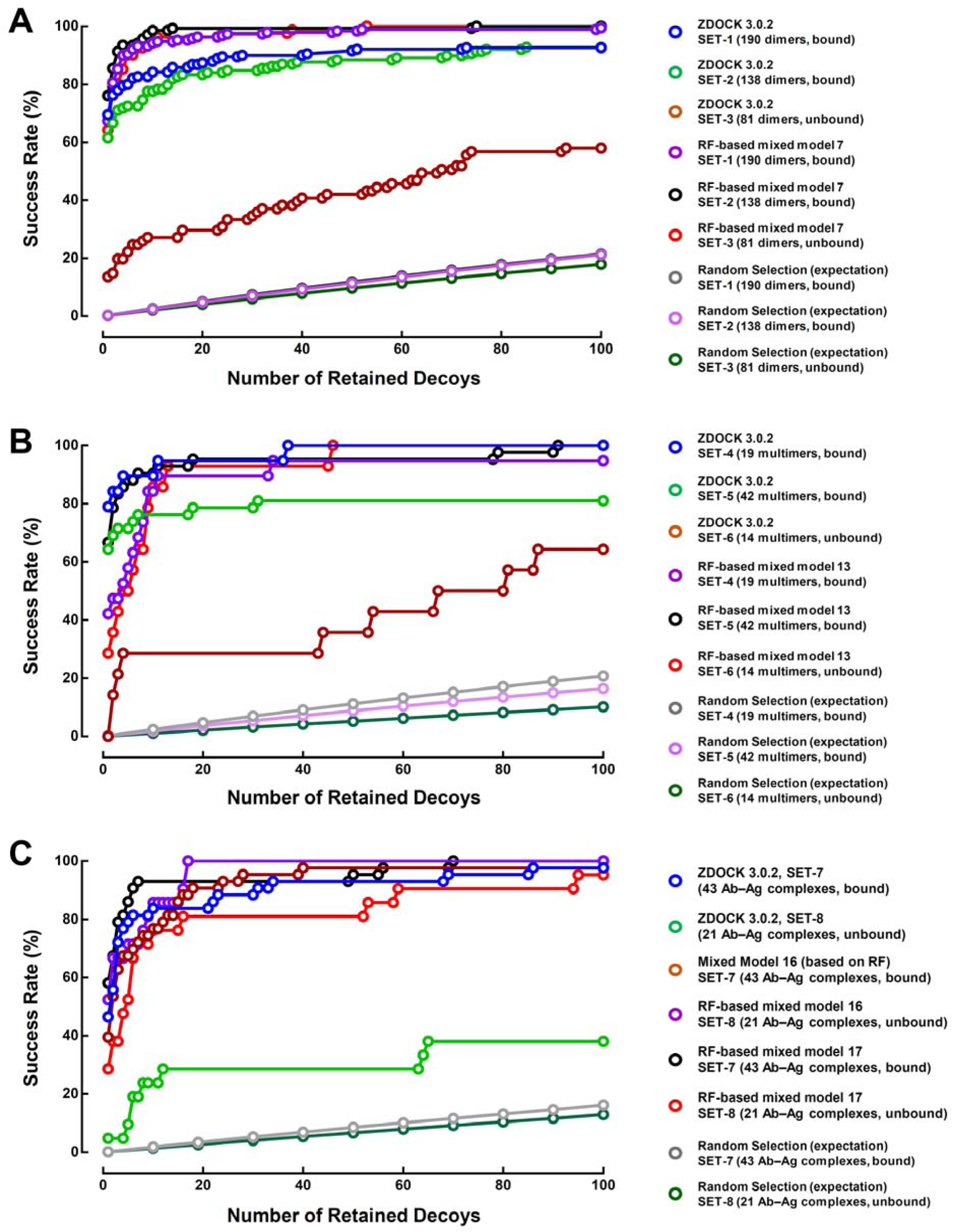
The performances of ZDOCK3.0.2 and several representative RF-based mixed models for protein–protein binding interface selection. **A.** The performances of ZDOCK3.0.2, RF-based mixed model 7 (**Table 7**) and random selection in the dimer group. **B.** The performances of ZDOCK3.0.2, RF-based mixed model 13 (**Table 7**) and random selection in the multimer group. **C.** The performances of ZDOCK3.0.2, RF-based mixed models 16 and 17 (**Table 7**) and random selection in the antibody-protein antigen group. The Ab–Ag complexes refer to the antibody–protein antigen complexes.

### 3.3. The apparent gap between affinity prediction and interface selection

Theoretically, a perfect predictive model should be able to calculate the binding affinity accurately and also correctly select the near-native interfaces/docking decoys at the same time. However, earlier studies have indicated that there is an apparent gap between the prediction of protein–protein binding interfaces (*i.e.*, structure prediction) and the prediction of protein–protein binding affinities [44,45]. This problem was explored in the present study. First, we determined the performances of previous area-based affinity predictive models in interface selection. The 97 area-based affinity predictive models developed recently by us [35,36] were employed to select near-native structures; likewise, the 3 new linear interface selection models (**Table 3**) and the 60 representative neural network, random forest and mixed models (**Tables 4**–**7**) were also used to predict the binding affinities of the protein–protein complexes.

Next, we also determined the performances of the new interface selection models in affinity prediction. Several representative affinity models were selected based on their performances for interface selection and affinity prediction (**Supplementary File 3**). While the affinity models 5, 30 and 94 have better interface selection performances than those of other models, the affinity models 58 and 60 possess better affinity predictive performances in all the affinity sets and subsets (**Fig. 4A, 4B, Supplementary File 4**). As shown in **Table 8**, the success rates of affinity model 5 for dimers in SET-1, SET-2 and SET-3 are higher than 20%; the success rates of affinity model 30 for multimers in SET-4, 5 and 6, affinity model 94 for the antibody–protein antigen complexes in SET-7 and 8, and affinity models 58 and 60 for the complexes in SET-3, SET-6 and SET-8 are above 10%. The Pearson’s correlation coefficient (*R*) for affinity model 5 in affinity SUBSET-1 (33 dimers) or affinity model 30 in affinity SUBSET-2 (9 multimers) or affinity model 94 in affinity SUBSET-3 (37 antibody-protein antigen complexes), is 0.76 or 0.69 or 0.54, respectively; and the *R* values of the affinity models 58 and 60 are all above 0.65 in the combined SET, SUBSET-1, SUBSET-2 and SUBSET-3 (**Fig. 4A, 4B, Supplementary File 4**). It is apparent that the better performances in affinity prediction do not always couple with better performances in interface selection. Overall, the performances of affinity models in interface selection are inferior to those models designed exclusively for interface selection.

**Figure 4.**
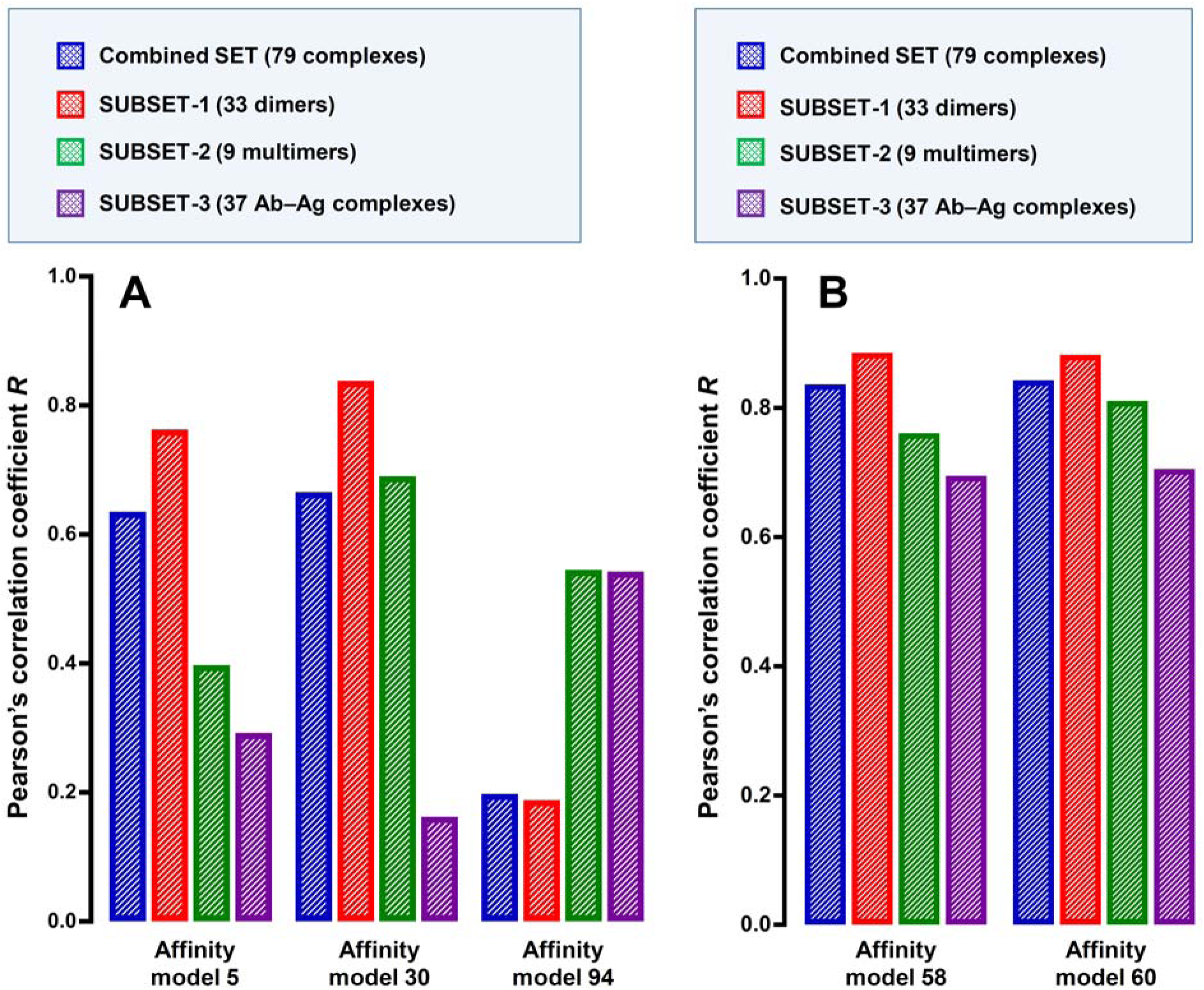
The performances of several previous affinity predictive models in binding affinity selection. **A.** The performances of the affinity models 5, 30 and 94. **B.** The performances of the affinity models 58 and 60. The performances of binding affinity prediction are represented by the Pearson’s correlation coefficient (*R*). The Ab–Ag complexes refer to the antibody–protein antigen complexes.

**Table 8.** Performances of the 5 previous affinity predictive models in binding interface selection.

| Model name and No. | Targeted applications for interface selection | Success rate (%) (top 10 decoys, L_RMSD < 5 Å) |  |  |  |  |  |  |  |
| --- | --- | --- | --- | --- | --- | --- | --- | --- | --- |
|  |  | SET-1<br>(190 dimers,<br>bound) | SET-2<br>(138 dimers,<br>bound) | SET-3<br>(81 dimers,<br>unbound) | SET-4<br>(19 multimers,<br>bound) | SET-5<br>(42 multimers,<br>bound) | SET-6<br>(14 multimers,<br>unbound) | SET-7 (43<br>Ab–Ag<br>complexes,<br>bound) | SET-8<br>(21 Ab–Ag<br>complexes,<br>unbound) |
| **Affinity model 5** | Dimer | 39.47 | 35.51 | 22.22 | 36.84 | 14.29 | 7.14 | 2.33 | 4.76 |
| **Affinity model 30** | Dimer and multimer | 32.63 | 26.09 | 14.81 | 31.58 | 11.90 | 14.29 | 6.98 | 0.00 |
| **Affinity model 58** | Dimer, multimer, and<br>Ab–Ag complex | 10.53 | 10.14 | 1.23 | 15.79 | 0.00 | 0.00 | 9.30 | 0.00 |
| **Affinity model 60** |  | 7.89 | 7.97 | 2.47 | 26.32 | 7.14 | 7.14 | 4.65 | 0.00 |
| **Affinity model 94** | Ab–Ag complex | 14.21 | 16.67 | 12.35 | 21.05 | 11.90 | 0.00 | 11.63 | 14.29 |

Note: The affinity model 5 is the linear model 5; the affinity model 30 is the generated nonlinear model (linear fitting) 7; the affinity model 58 is the NN-based mixed model 1; the affinity model 60 is the NN-based mixed model 3 for protein–protein binding affinity prediction [31]; and the affinity model 94 is the RF-based mixed model 1 for antibody–protein antigen affinity prediction [32]. The Ab–Ag complexes refer to the antibody–protein antigen complexes.

In this study, the performances of 63 models for binding interface selection were also studied for their affinity prediction ability by using the affinity datasets (**Supplementary File 5**). The highest *R* values are 0.48 in the combined SET (linear model 3), 0.66 in its SUBSET-1 (RF model 3), 0.88 in its SUBSET-2 (NN-based mixed model 5), and 0.37 in its SUBSET-3 (linear model 3). The interface selection performance of linear model 3 for dimers and multimers is not outstanding. The better interface selection performances usually do not imply better predictive powers for affinity. Overall, the best performances of the area-based models for near-native structure selection in affinity prediction are not as good as those of the best models exclusively designed for affinity prediction.

## 4. DISCUSSION

The present study seeks to make a tentative exploration of the protein–protein binding interface selection problem from an area perspective rather than by using the traditional computational approaches. The results gathered in this study illustrate the potential ability of the area-based methods. According to the performances of the linear, neural network, random forest and mixed models, this study suggests the potential existence of area-based pairing rules for protein–protein interactions. It is hopeful that when the area-based models are more refined to reach a higher level of effectiveness and universality, the theoretical basis might slowly emerge in the future.

In practice, a function or equation can be treated as the paring rules for proteinLprotein interactions if it can discriminate the near-native interfaces/docking decoys from the non-native ones with a high success rate. In this work, the explicit and implicit functions for the quantitative area–L_RMSD relationship are explored using linear regression and machine leaning methods to select the near-native interfaces. The signs (+ or –) of the coefficients of the shared descriptors in linear models 1–3 (**Table 3**) are: **– – +** for ***A*_5_** (*LSA* of basic *AA*s), **– – –** for ***A*_10_** (nonpolar *AA*s∼nonpolar *AA*s), **+ + +** for ***A*_12_** (acidic *AA*s∼acidic *AA*s), **– – –** for ***A*_14_** (basic *AA*s∼polar *AA*s), **– – –** for ***A*_15_** (basic *AA*s∼acidic *AA*s), **+ + +** for ***A*_17_** (nonpolar *AA*s∼acidic *AA*s) and **– – –** for ***A*_18_** (polar *AA*s∼acidic *AA*s), respectively. While the higher values of the ***A*_10_**, ***A*_14_**, ***A*_15_** and ***A*_18_** are associated with the lower L_RMSD values for the dimers, multimers and antibody-protein antigen complexes, the lower values of the ***A*_12_** and ***A*_17_** are associated with lower L_RMSD values for the dimers, multimers and antibody-protein antigen complexes. In addition, the higher ***A*_5_** values are accompanied with lower L_RMSD values for the dimers and multimers but higher L_RMSD values for antibody-protein antigen complexes. It is apparent that different kinds of areas make different contributions in interface selection. The geometric shape complementary, the attractive and repulsive energies, and the favorable and unfavorable protein-solvent interactions are all implied in the different kinds of interface and surface areas, which determine the overall reasonability of the conformations with different binding regions and orientations. If the binding regions between the two binding partners are unique, then their quantitative area–L_RMSD relationship around the binding regions likely should exist; if not, the relationship may be ambiguous as the L_RMSD for a given conformation likely is not unique. Therefore, the uniqueness of the binding regions between the two binding partners should also be considered when predicting the structures of the protein–protein complexes.

Overall, the performances of the single metric, *i.e.*, the area-based linear functions among the interfaces/decoys generated by docking the unbound proteins are better than the performances of ZDOCK3.0.2 which is based on geometric shape complementarity, electrostatic energy and knowledge-based pair potentials [23,34]. The knowledge-based pair potentials are derived based on the numbers of different amino acid types and the pairs in the structures of proteins and protein-protein complexes [22]. It may be more suitable to explore the rules of interface specificity from a geometric area perspective than from the numbers of different amino acid types and pairs. We find that the RF-based mixed models achieve high success rates for dimers (>90%), multimers (>80%) and antibody-protein antigen complexes (>75%) (**Table 7**). The area-based models open a new window for exploring and understanding the principles of protein–protein interactions. Parallel to the base-pairing rules for understanding the structures of DNA molecules, the area-based methods shed lights on the pairing rules for protein–protein interactions and also on the protein–protein binding mechanisms.

Based on the results of the area-based models for affinity prediction and interface selection, we notice that the gap between protein–protein binding interface selection (*i.e.*, structure prediction) and affinity prediction does exist. While structure prediction mostly focuses on the geometric feasibility and reasonability, affinity prediction gives special importance to energy reasonability. In theory, the native structures should be the geometric conformations with the lowest binding energy. In computational practice, only using the energy cannot always select the near-native structures successfully [13]. The experimental binding affinity or binding energy in protein–protein interactions is not entirely consistent with the theoretically-calculated binding energy based on the native structures of the protein–protein complexes [35,36,43,46]. Even if we employ more rigorous methods such as MM/PBSA and MM/GBSA (in need of more time and computational resources), the results of affinity calculation and interface selection are often still not ideal [47]. While the experimental determination of the binding affinity usually takes into consideration the complete binding process, the native structures of the protein–protein complex often are just snapshots during the process, which may only partially reflect the experimental binding affinity. The two binding partners in the native structures of the transient protein–protein complexes often would dissociate from each other during biological processes. The balance between association and dissociation of the protein binding partners requires relative stability of the binding interface and the assistance of the surface-solvent interactions. A better understanding of the principles and problems intrinsically associated with affinity prediction and interface selection may aid in the development of more effective prediction methods in the future.

It should be noted that a potential problem of the area-based models relates to the non-commutativity of the receptor and ligand surface areas. The linear area-based empirical models are very superficial and the results often are not good enough, which is as expected. The more advanced nonlinear explicit formations which have better performances and commutativity of the receptor and ligand surface areas may be developed in the future. Another potential problem relates to the lack of consideration of the conformational changes (flexibility) upon partner’s binding for the rigid body-based docking process which relies on the partners’ structures at the unbound state. It is expected that when sufficient computational resources become available, molecular dynamics simulation may be effectively employed to generate more conformations with sufficient consideration of the structural flexibility [48]. The area–flexibility relationship in protein structures has been explored in previous studies [49,50]. Incorporating the area–flexibility relationship into docking and interface selections may enhance the current predictive power for the structures of the protein–protein complexes.

In this study, the success rates for the top 10 decoys with L_RMSD < 5 Å of the mixed models without explicit formations have achieved high values (>90% for the dimer group, >80% for the multimer group, and >75% for the antibody–protein antigen group), which clearly indicates the effectiveness of the area-base methods in interface selection. Given the disparities in the performances of a selected area-based model for interface selection in different groups of datasets (as listed in **Table 1**), it is essential to consider the dimers, multimers, and antibody–protein antigen complexes separately in order to achieve higher levels of performance. Additionally, the contributions of non-interface features to protein–protein interactions (binding energy/affinity in protein–protein complex and geometric structure of protein–protein complex) have been explored in several previous works in recent years [35,36,43,46,51–54]. It is also worth mentioning that there are still rooms for improvements in the performances of the area-based models with explicit formations for both interface selection (*i.e.*, structure prediction) and affinity prediction. With improved size and quality of the protein datasets (*e.g.*, with tens of thousands of high-quality protein–protein complexes), it is expected that individual consideration of the 20 amino acid types may further improve the performances of the area-based interface selection approach. The interface-specific and global surface contributions will be analyzed and discussed using the better models in the future. The emphasis of our present work is to demonstrate that the area-based models alone indeed can achieve comparable or even better performances as the combinational scoring functions of ZDOCK3.0.2 for the docking of unbound structures, which is rather remarkable. The best area-based models in this work jointly employ the interface and surface features. According to the performances, the RF-based mixed model 7 is recommended for interface selection of dimers; the RF-based mixed model 13 for interface selection of multimers; and the RF-based mixed models 16 and 17 for interface selection of antibody–protein antigen complexes.

It is of note that while some of the big AI models developed in recent years, such as AlphaFold-Multimer [55] and AlphaFold3 [56,57], can provide predicted structures of protein–protein complexes, the accuracy of the structural prediction for the protein–protein complexes is still a key scientific problem which is presently not fully resolved. According to the results of CASP16, AI models can generate the protein–protein binding interfaces lacking co-evolutionary signals [58]. The corresponding near-native structures of such protein–protein complexes can be generated by the protein–protein docking methods [7]. Additionally, selecting the best structural models from a large numbers of possible decoys is still a major bottleneck [58]. The present work has developed new selection methods for the structural models of protein–protein complexes from an area perspective, which may aid in the future development of methods for more accurately predicting the structures of the protein–protein complexes.

## 5. CONCLUSION

The pairing rules across protein–protein binding interface are important for structure prediction of the protein–protein complexes. In this study, the pairing rules are explored from a geometric area perspective. The predictive powers of the linear and nonlinear area-based models developed in this study for interface selection reveal the possible existence of the area-based pairing rules for protein–protein interactions. In the future, more exhaustive explorations of the paring rules by employing larger sets of higher quality structures of the protein–protein complexes may help better refine the pairing rules developed in this study and reach more precise conclusions on the pairing rules.

## Data availability statement

All the data of the study are available upon request.

## Conflict of interest

The first author has a pending patent application related to this work.

## Author Contributions

Conceptualization, Y.X.Y.; B.T.Z.; methodology, Y.X.Y.; formal analysis, Y.X.Y., B.T.Z.; investigation, Y.X.Y., B.T.Z.; resources, B.T.Z.; data curation, Y.X.Y.; writing—original draft preparation, Y.X.Y., B.T.Z.; writing—review and editing, Y.X.Y., B.T.Z.; visualization, Y.X.Y.; resource maintenance: Y.X.Y.; supervision, B.T.Z.; project administration, B.T.Z.; funding acquisition, B.T.Z.

## Acknowledgements

This work is supported by research grants from Shenzhen Key Laboratory of Steroid Drug Discovery and Development (No. ZDSYS20190902093417963), Shenzhen Peacock Plan (No. KQTD2016053117035204) and Shenzhen Bay Laboratory (No. SZBL2019062801007).

## Notes

This work was supported, in part, by research grants from Shenzhen Peacock Plan (No. KQTD2016053117035204), Shenzhen Key Laboratory of Steroid Drug Discovery and Development Research (No. ZDSYS20190902093417963), Shenzhen Bay Laboratory (No. SZBL2019062801007), 2022 Stable Funding Support Program for Shenzhen Institutions of High Learning, and Longgang District Science and Technology Bureau’s Key Laboratory Program.

### Competing Interest Statement

The authors have declared no competing interest.

